# Shared and Divergent Features of Cardiac Transcriptome and Glucose Metabolism Markers in Human and Mouse HFpEF

**DOI:** 10.64898/2026.08.20.746109

**Authors:** Kajol Thapa, Kleio-Maria Verrou, Erjola Rapushi, Georgios Siokatas, Karthickeyan Chella Krishnan, Nike Bharucha, Brendan J Keating, Markus Meyer, Ioannis Karakikes, Konstantinos Drosatos

## Abstract

Heart Failure with Preserved Ejection Fraction (HFpEF) is more prevalent in females and is associated with altered cardiac glucose metabolism. However, whether these metabolic alterations are conserved across sexes and between humans and widely used cardiometabolic mouse model of HFpEF remains unclear. We investigated species-, sex-, and ventricle-specific conserved and divergent features of HFpEF.

Cardiometabolic HFpEF was induced in mice using the “two-hit” model (high-fat diet + L-NAME), followed by assessment of cardiac function, RNA sequencing, and protein expression in the right (RV) and left (LV) ventricles. Published human HFpEF RV and LV RNA-seq datasets were reanalyzed and compared with our mouse data.

Only male HFpEF mice recapitulated human phenotype of increased RV GLUT1 protein. In contrast, mouse GLUT1 was downregulated in RV of females and in the LV of both sexes, whereas GLUT4 protein remained unchanged. Cardiac PDK4 transcript and protein levels increased in the RV and LV of mice. Conversely, human PDK4 mRNA levels were reduced in the RV with HFpEF and unchanged in LV. Cardiac transcriptome analysis in mice revealed extensive alterations in LV, particularly in females, with enrichment of inflammatory pathways. Cross-species analysis demonstrated greater conservation of HFpEF-associated signatures in the RV than the LV. Furthermore, number of differentially expressed transcripts in human LV increased substantially after excluding patients with atrial fibrillation or diabetes.

Overall, the RV of the “two-hit” model more closely resembles human HFpEF. The cardiac transcriptome reflects sexual dimorphism, and conserved signatures are primarily associated with metabolic alteration, mitochondrial dysfunction, and cellular stress.

**HIGHLIGHTS:**

- The human RV GLUT1 phenotype is reproduced in male mice with HFpEF.
- HFpEF-related cardiac transcriptome changes are more pronounced in female mice.
- RV shows greater cross-species conservation than LV in HFpEF.
- Conserved transcriptomic changes primarily reflect alterations in metabolism and cellular stress.
- Humans and mice with HFpEF have opposite PDK4 expression profiles.

**GRAPHICAL ABSTRACT:** 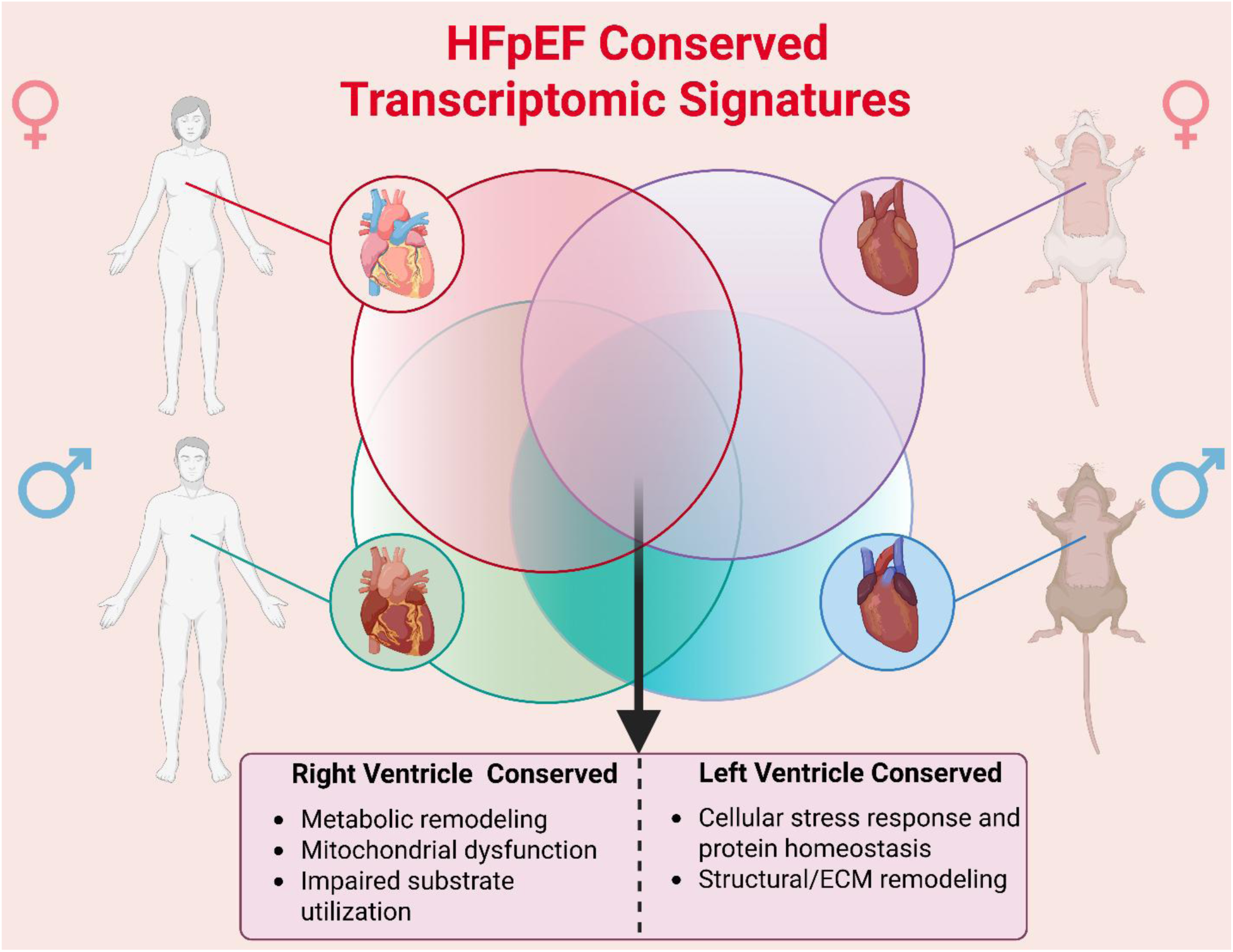

## 1. Introduction

Heart failure (HF) is a heterogeneous syndrome affecting approximately 64 million individuals worldwide and characterized by inadequate cardiac output[1]. Based on left ventricular (LV) ejection fraction (LVEF), HF is categorized into reduced (HFrEF), mildly reduced (HFmrEF), and preserved (HFpEF) phenotypes[2]. HFpEF accounts for about half of all HF cases [2] and has increasingly become a focus of clinical and preclinical research.

Obesity, diabetes, hypertension, and aging are primary risk factors for HFpEF, underscoring the critical role of metabolic dysregulation in disease progression[3]. A healthy adult heart relies primarily on fatty acid oxidation for ATP synthesis. This metabolic profile is often compromised in HF, which is often accompanied by a shift towards glucose utilization [4, 5]. Clinical studies have shown that glucose-lowering medications, such as sodium-glucose cotransporter 2 (SGLT2) inhibitors and glucagon-like peptide-1 (GLP-1) receptor agonists, improve cardiovascular outcomes in patients with HFpEF, thus emphasizing the importance of systemic glucose dysregulation in HFpEF pathophysiology[6, 7]. On the other hand, findings on the regulation of glucose metabolism are conflicting. A recent study that analyzed right ventricles (RV) isolated from patients with HFpEF reported increased protein levels of Glucose Transporter (GLUT)1 [8]. The same study found lower abundance of glucose-derived metabolites. Conversely, another study found decreased *SLC2A1* (GLUT1) transcript levels[9]. Studies in animal models have not helped to resolve these controversies. In particular, recent murine HFpEF models have shown reduced *Slc2a1* mRNA expression, highlighting inconsistencies across species [10, 11]. This discrepancy, combined with the incomplete understanding of sex- and chamber-specific metabolic responses, represents a critical gap in elucidating HFpEF pathophysiology.

This study investigated the regulation of glucose metabolism-related gene expression in the context of sex- and cardiac compartment interspecies and intraspecies similarities and differences. To this end, we used the cardiometabolic “two-hit” HFpEF mouse model, which involves treating wild-type mice with a high-fat diet (HFD) and Nω-nitro-L-arginine methyl ester (L-NAME) [12]. For both early- and late-phase HFpEF, we compared RV and LV expression of GLUT1, GLUT4, and Pyruvate Dehydrogenase Kinase 4 (PDK4) in mice. In late-phase HFpEF, we also compared murine RV and LV cardiac transcriptomes with published human HFpEF RNA-seq datasets.[13, 14]. Our analyses revealed distinct transcriptomic profiles and conserved transcript changes shared across species, sexes, and chambers, highlighting core signatures that drive myocardial dysfunction in HFpEF. Therefore, our study suggests novel interspecies similarities that may enhance bench-to-bedside translatability.

## 2. Methodology

### 2.1. Animals

All experimental procedures were conducted in accordance with the National Institutes of Health Guide for the Care and Use of Laboratory Animals and were approved by the IACUC at the University of Cincinnati. Eight-week-old male C57BL/6N mice (The Jackson Laboratory; Strain: 005304) were assigned to two groups. The HFpEF group received a high-fat diet (HFD; rodent diet with 60 kcal% fat) and L-NAME (0.5 g/L, pH=7.4) in the drinking water for 5 or 12 weeks. The control group was fed a standard chow diet with regular drinking water for the same duration. Similarly, eight-week-old female C57BL/6J mice were fed an HFpEF diet for 12 weeks. Cardiac function and blood pressure were assessed using echocardiography and the non-invasive tail-cuff method, respectively.

### 2.2. Echocardiography

Echocardiography was performed on HFpEF and control mice at weeks 5 and 12 after treatment using the Vevo F2 ultrasound system (VisualSonics), as previously described [15]. Mice were anesthetized with 2–3% isoflurane and maintained at 0.5–1% during imaging. After hair removal and application of ultrasound gel, mice were positioned on a heated platform for image acquisition. M-mode and B-mode imaging were used to assess cardiac structure and function. Ejection fraction was measured from short-axis M-mode images. Pulse-wave Doppler was used to determine peak early diastolic transmitral flow velocity (E wave), and tissue Doppler imaging was used to measure early diastolic myocardial relaxation velocity (e′). The E/e′ ratio was calculated as an index of left ventricular filling pressure and diastolic function. Heart rate and body temperature were continuously monitored, and all images were analyzed using Vevo Lab software.

### 2.3 Non-Invasive Blood Pressure Monitoring

Systolic, mean, and diastolic blood pressures were measured non-invasively in conscious mice using the BP-2000 tail-cuff system (Visitech Systems). Mice were acclimated to the restrainer and the measurement procedure for 5 consecutive days prior to randomization, and blood pressure was recorded on the 6th day. During measurements, mice were placed in restrainers on a warming platform maintained at 39°C for at least 10 minutes to promote tail vasodilation. Thirty measurements were obtained for each mouse at each time point, and values were averaged to determine the final blood pressure.

### 2.4 RNA extraction, purification, and gene expression

Cardiac tissue (both RV and LV) was dissolved in 1 mL TRIzol lysis reagents (ThermoFisher, 15596026) as described previously [15]. For cDNA synthesis, DNase I, Amplification Grade was used to digest single- and double-stranded DNA. The High-Capacity cDNA Reverse Transcription Kit (Applied Biosystems 4368814) was then used to convert RNA into single-stranded cDNA, and finally RT-PCR was performed using StepOnePlus Real-Time PCR system and SYBR Green master mix (Applied Biosystems, 4472903). Expression of the targeted gene was then normalized to 36B4 housekeeping gene. Relative quantification of a targeted gene was performed using the 2^−ΔΔCt^ method. The sequences of all primers used in this study are listed in Supplementary Table 1.

### 2.5 RNA-seq Analysis

Total RNA was extracted from ventricular tissue as described above. RNA quantity and purity were assessed using a NanoDrop spectrophotometer (Thermo Fisher Scientific), and RNA integrity was evaluated using an Agilent 2100 Bioanalyzer (Agilent Technologies). Samples with RNA Integrity Number (RIN) ≥7 were used for library preparation. RNA-seq libraries were prepared using the NEBNext Ultra II RNA Library Prep Kit (New England Biolabs) according to the manufacturer’s instructions. LV samples were sequenced as paired-end 150-bp reads on an Illumina NovaSeq X platform, whereas RV samples were sequenced as single-end 94-bp reads on an Illumina NovaSeq X Plus platform. The average sequencing depth was ∼24 million reads per sample.

Raw reads were aligned to the human (hg38/UCSC) or mouse (mm39/GRCm39) genome using STAR, and gene-level counts were generated with featureCounts (v2.0.0 and 2.0.6) using paired-end settings (-p), same-chromosome and properly paired read filters (-B, -C; human only), exon features (-t exon), gene_id annotation (-g gene_id), a minimum overlap of 10 bp (--minOverlap 10; human only), and fragment length limits of 30–600 bp (-d 30 -D 600; human only). Human samples used UCSC hg38 annotations, whereas mouse samples used GENCODE vM37 (mm39) or Ensembl GRCm39.112 annotations.

Differential expression analysis was performed in R (v4.4.3) using DESeq2 (v1.46.0). Gene annotations were obtained from Ensembl BioMart, and only protein-coding genes were retained. Genes with <5 counts in <25% of samples per group were excluded. Data were normalized in DESeq2, with differential expression assessed using the design formula ∼ sex + condition or ∼ condition for sex-stratified analyses. Variance-stabilizing transformation (VST) was used for PCA and visualization. DEGs were defined as adjusted *P*<0.05 using the Benjamini–Hochberg correction. Gene set enrichment analysis was performed using clusterProfiler, and visualizations were generated with ggplot2, pheatmap, and DESeq2.

### 2.6 Protein expression analysis

Myocardial tissue was homogenized in RIPA buffer supplemented with protease and phosphatase inhibitors (15 μL/mg tissue; Thermo Fisher Scientific, A32961) and incubated on ice for 30 min before centrifugation at 12,000 × *g* for 15 min at 4°C. Protein concentration was determined using a BCA assay. For Western blot analysis, 60 μg of protein was used for GLUT1 and GLUT4, and 30 μg for PDK4. Samples were prepared using optimized conditions (GLUT1, unheated; GLUT4, 45°C for 15 min; PDK4, 90°C for 5 min), separated by 10–12% SDS-PAGE, and transferred to PVDF membranes using a Turbo Transfer system. Membranes were blocked with 5% BSA and incubated overnight at 4°C with primary antibodies against GLUT1 (Cell Signaling, 12939S), GLUT4 (Proteintech, 66846-1-Ig), PDK4 (Abcam), and β-actin (Santa Cruz Biotechnology, sc-47778). After incubation with secondary antibodies (1:5000), protein bands were visualized using a LI-COR Odyssey CLx Infrared Imager and quantified in ImageJ after normalization to β-actin.

### 2.7 Statistical Analysis

Statistical analysis was performed using GraphPad Prism (Graph Pad Software version 11.0.1) and R-version 4.2.1. All the data (N≥ 5) were presented as mean± standard error mean (SEM). First, data was checked for normality using the Shapiro-Wilk test (P<0.05). Data that passed the normality test (P>0.05) were subjected to statistical analysis, including two sample-t tests or one-way ANOVA when comparing two groups or more than two groups respectively. If the data did not pass the normality test, they were subjected to either log-transfer or Mann-Whitney and Kruskal-Wallis non-parametric test. The presence of outliers was confirmed using the GraphPad prism or manually by calculating the interval (mean ± 2*standard deviation). P-value less than 0.05 was considered statistically significant.

### 2.8 Data Availability

The bulk RNA-sequencing datasets generated in this study have been deposited in the Gene Expression Omnibus under accession numbers **GSE338090** (right ventricles) and **GSE337935** (left ventricle). All other supplementary data generated during this study have been deposited in Mendeley and are available under the reserved DOI: **10.17632/r5n5x98tf5.1**.

## 3. Results

### 3.1 Early and sustained HFpEF phenotype in male and female mice following HFD/L-NAME treatment

As previously described[12], eight-week-old female C57BL/6J and male C57BL/6N mice were subjected to a high-fat diet (HFD) supplemented with L-NAME in the drinking water (HFpEF diet) for either 5 weeks (early stage) or 12 weeks (late stage). Age-and strain-matched control mice received standard chow and regular drinking water for the same durations. Mice were randomly assigned to experimental groups. HFpEF diet-fed female mice exhibited a significant increase in body weight at both the early (Figure 1A) and late stage (Figure 1B), which was accompanied by impaired glucose tolerance (Figure 1C-F), compared to chow-fed control mice. Accordingly, both systolic and diastolic blood pressure were increased at both time points (Fig. 1G, 1H). In addition, these mice developed cardiac hypertrophy (Figure 1I) and pulmonary edema (Figure 1J). Echocardiographic analysis showed that, as expected in HFpEF, left ventricular ejection fraction remained unchanged (Figure 1K). Diastolic dysfunction was evident at 5 weeks and persisted at 12 weeks, as indicated by an increased E/e′ ratio (Figure 1L). Likewise, male C57BL/6N mice that were subjected to the same HFpEF diet displayed the same phenotype in body weight (Figure 1M, 1N), glucose intolerance (Figure 1O–R) systolic (Figure 1S) and diastolic blood pressure (Figure 1T), cardiac hypertrophy (Figure 1U), pulmonary edema (Figure 1V), preserved ejection fraction (Figure 1W), and diastolic dysfunction, as evidenced by an increased e/e′ ratio (Figure 1X), compared to chow-fed control mice. Notably, the severity of diastolic dysfunction appeared more pronounced in male mice compared to females (Figure 1L, 1X).

**Figure 1.**
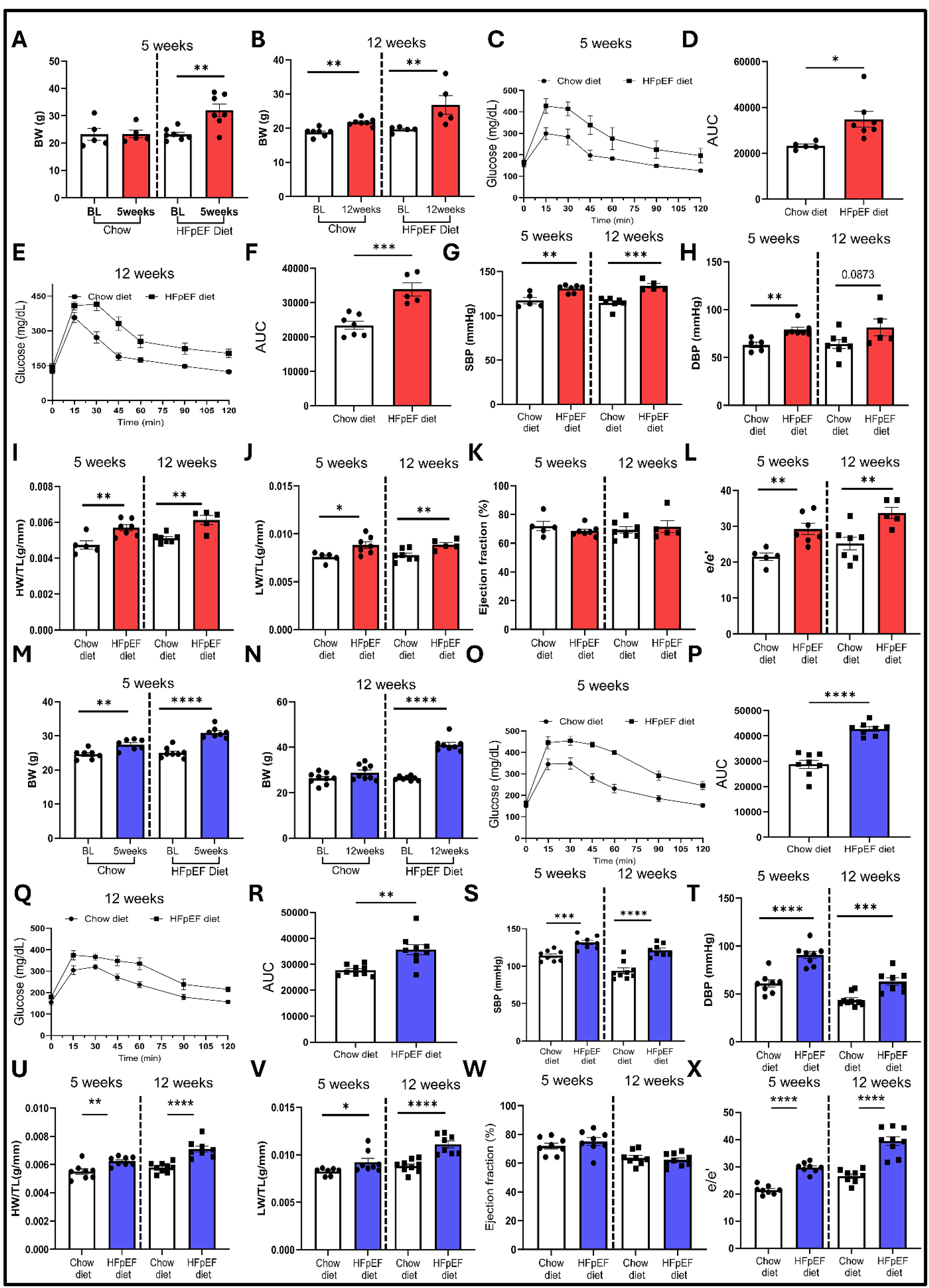
Establishment and characterization of a diet-induced HFpEF model in female and male mice. (A) Body weight (BW) of female mice at 5 weeks compared to baseline. (B) Body weight (BW) of female mice at 12 weeks compared to baseline. (C) Glucose tolerance test (GTT) in female mice at 5 weeks. (D) Area under the curve (AUC) of the GTT in female mice at 5 weeks. (E) Glucose tolerance test (GTT) in female mice at 12 weeks. (F) Area under the curve (AUC) of the GTT in female mice at 12 weeks. (G) Systolic blood pressure (SBP) in female mice. (H) Diastolic blood pressure (DBP) in female mice. (I) Heart weight-to-tibia length ratio (HW/TL) in female mice. (J) Lung weight-to-tibia length ratio (LW/TL) in female mice. (K) Left ventricular ejection fraction (EF) in female mice. (L) Ratio of early diastolic transmitral flow velocity to early diastolic mitral annular velocity (e/e′) in female mice. (M) Body weight (BW) of male mice at 5 weeks compared to baseline. (N) Body weight (BW) of male mice at 12 weeks compared to baseline. (O) Glucose tolerance test (GTT) in male mice at 5 weeks. (P) Area under the curve (AUC) of the GTT in male mice at 5 weeks. (Q) Glucose tolerance test (GTT) in male mice at 12 weeks. (R) Area under the curve (AUC) of the GTT in male mice at 12 weeks. (S) Systolic blood pressure (SBP) in male mice. (T) Diastolic blood pressure (DBP) in male mice. (U) Heart weight-to-tibia length ratio (HW/TL) in male mice. (V) Lung weight-to-tibia length ratio (LW/TL) in male mice. (W) Left ventricular ejection fraction (EF) in male mice. (X) Ratio of early diastolic transmitral flow velocity to early diastolic mitral annular velocity (E/e′) in male mice. Statistical analyses were performed using GraphPad Prism (version 11.0.1). Comparisons between two groups were performed using an unpaired, two-tailed Student’s *t*-test. Statistical significance is indicated as *P* < 0.05, P < 0.01, *P* < 0.001, and P < 0.0001. Abbreviations: BW, body weight; SBP, systolic blood pressure; DBP, diastolic blood pressure; HW/TL, heart weight-to-tibia length ratio; LW/TL, lung weight-to-tibia length ratio; GTT, glucose tolerance test; AUC, area under the curve; EF, ejection fraction; E/e′, ratio of early diastolic transmitral flow velocity (E) to early diastolic mitral annular velocity (e′).

### 3.2 The male mouse RV partially recapitulates human HFpEF-associated glucose metabolism markers

Driven by a human study that revealed higher cardiac GLUT1 protein expression in the RV septum of patients with HFpEF[8], we assessed its expression in the RV of mice with HFpEF and chow-fed control animals at the early (5 weeks) and late (12 weeks) stages of the disease. In contrast to the observation in humans, the RV of female mice with HFpEF showed a consistent reduction in GLUT1 protein (Figure 2A-C) and mRNA (Figure 2D) levels at both the early and late stages. On the other hand, GLUT1 expression in male mice was consistent with the human phenotype, showing a significant increase in GLUT1 protein (Figure 2E-G) despite the reduction of GLUT1 mRNA at 5 weeks and a 40% suppression at 12 weeks (Figure 2H).

**Figure 2.**
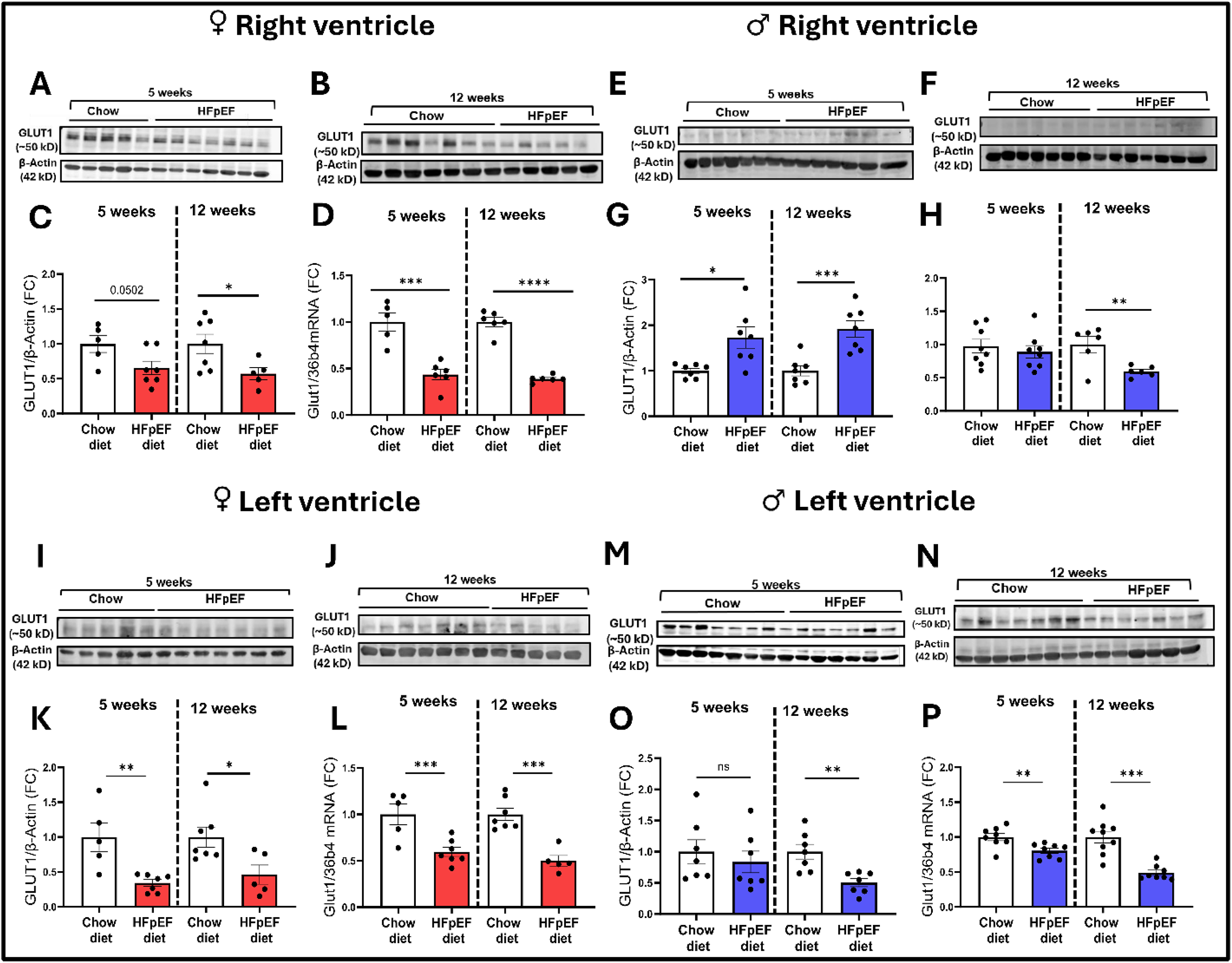
Sex- and chamber-specific regulation of GLUT1 in HFpEF. Mice were assigned to either an HFpEF group (high-fat diet plus L-NAME in drinking water; *n* = 5–7 per sex) or a chow-fed control group (*n* = 5–7 per sex). Right ventricular (RV) and left ventricular (LV) tissues were collected after 5 weeks (early phase) and 12 weeks (late phase). GLUT1 protein abundance was assessed by immunoblotting with densitometric quantification normalized to β-actin, and GLUT1 mRNA expression was normalized to 36B4. (A) Immunoblots of GLUT1 protein expression in the RV of female mice at 5 weeks. (B) Immunoblots of GLUT1 protein expression in the RV of female mice at 12 weeks. (C) Quantification of GLUT1 protein abundance in the RV of female mice at 5 and 12 weeks. (D) GLUT1 mRNA expression in the RV of female mice at 5 and 12 weeks. (E) Immunoblots of GLUT1 protein expression in the RV of male mice at 5 weeks. (F) Immunoblots of GLUT1 protein expression in the RV of male mice at 12 weeks. (G) Quantification of GLUT1 protein abundance in the RV of male mice at 5 and 12 weeks. (H) GLUT1 mRNA expression in the RV of male mice at 5 and 12 weeks. (I) Immunoblots of GLUT1 protein expression in the LV of female mice at 5 weeks. (J) Immunoblots of GLUT1 protein expression in the LV of female mice at 12 weeks. (K) Quantification of GLUT1 protein abundance in the LV of female mice at 5 and 12 weeks. (L) GLUT1 mRNA expression in the LV of female mice at 5 and 12 weeks. (M) Immunoblots of GLUT1 protein expression in the LV of male mice at 5 weeks. (N) Immunoblots of GLUT1 protein expression in the LV of male mice at 12 weeks. (O) Quantification of GLUT1 protein abundance in the LV of male mice at 5 and 12 weeks. (P) GLUT1 mRNA expression in the LV of male mice at 5 and 12 weeks. Statistical analyses were performed using GraphPad Prism (version 11.0.1). Comparisons between two groups were performed using an unpaired, two-tailed Student’s *t*-test. Statistical significance is indicated as *P* < 0.05, P < 0.01, *P* < 0.001, and P < 0.0001. Abbreviations: GLUT1, glucose transporter 1; RV, right ventricle; LV, left ventricle.

GLUT1 expression in the LV showed a more consistent expression profile across sexes. Both female (Figure 2I-L) and male (Figure 2M-P) mice with HFpEF had lower GLUT1 mRNA levels at both the early and late stages of the disease. Notably, although GLUT1 mRNA expression was reduced in the LV of male mice at both 5 and 12 weeks, a corresponding decrease in GLUT1 protein levels was observed as a trend at 5 weeks and clearly at 12 weeks (Figure 2P). Collectively, these findings reveal a sex-, chamber-, and disease-stage-specific expression pattern of cardiac GLUT1 in HFpEF, with RV GLUT1 in male mice of the “two-hit” HFpEF model similar to that in human HFpEF patients.

To determine the effect of HFpEF on broader cardiac glucose metabolism beyond GLUT1, we also assessed the expression of GLUT4, the main insulin-responsive glucose transporter in the adult heart, which remains unaltered in the RV of humans with HFpEF[8]. The GLUT4 expression profile showed a notable discordance between mRNA and protein levels. While GLUT4 protein levels did not change (Figure 3A-C), mRNA levels were significantly reduced (Figure 3D) in the RV of female mice with HFpEF at both early and late disease stages. The same pattern of expression for GLUT4 protein (Figure 3E-G) and mRNA (Figure 3H) levels was observed in the RV of male mice with HFpEF. The same analysis for GLUT4 expression in the LV of the HFpEF mouse model showed similar trends. Specifically, female mice with HFpEF had unaltered protein levels (Figure 3I-K) but lower cardiac mRNA levels compared to chow-fed control mice at both early and late stages (Figure 3L). Male mice showed no change in protein levels (Figure 3M-O), and GLUT4 mRNA did not change, except for a minor suppression of LV GLUT4 mRNA at the early stage (Figure 3P).

**Figure 3.**
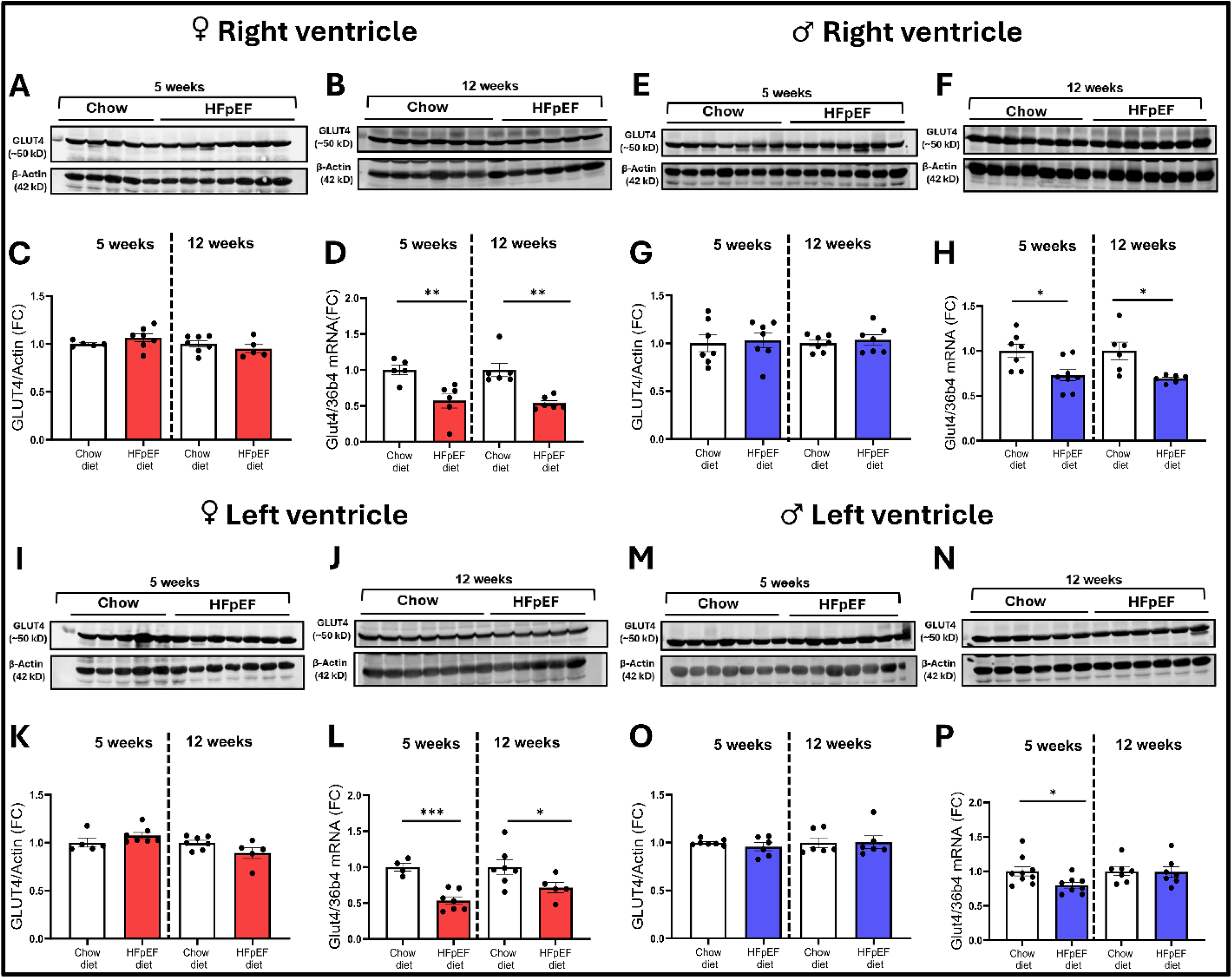
Sex- and chamber-specific regulation of GLUT4 in HFpEF. Mice were assigned to either an HFpEF group (high-fat diet plus L-NAME in drinking water; *n* = 5–7 per sex) or a chow-fed control group (*n* = 5–7 per sex). Right ventricular (RV) and left ventricular (LV) tissues were collected after 5 weeks (early phase) and 12 weeks (late phase). GLUT4 protein abundance was assessed by immunoblotting with densitometric quantification normalized to β-actin, and GLUT4 mRNA expression was normalized to 36B4. (A) Immunoblots of GLUT4 protein expression in the RV of female mice at 5 weeks. (B) Immunoblots of GLUT4 protein expression in the RV of female mice at 12 weeks. (C) Quantification of GLUT4 protein abundance in the RV of female mice at 5 and 12 weeks. (D) GLUT4 mRNA expression in the RV of female mice at 5 and 12 weeks. (E) Immunoblots of GLUT4 protein expression in the RV of male mice at 5 weeks. (F) Immunoblots of GLUT4 protein expression in the RV of male mice at 12 weeks. (G) Quantification of GLUT4 protein abundance in the RV of male mice at 5 and 12 weeks. (H) GLUT4 mRNA expression in the RV of male mice at 5 and 12 weeks. (I) Immunoblots of GLUT4 protein expression in the LV of female mice at 5 weeks. (J) Immunoblots of GLUT4 protein expression in the LV of female mice at 12 weeks. (K) Quantification of GLUT4 protein abundance in the LV of female mice at 5 and 12 weeks. (L) GLUT4 mRNA expression in the LV of female mice at 5 and 12 weeks. (M) Immunoblots of GLUT4 protein expression in the LV of male mice at 5 weeks. (N) Immunoblots of GLUT4 protein expression in the LV of male mice at 12 weeks. (O) Quantification of GLUT4 protein abundance in the LV of male mice at 5 and 12 weeks. (P) GLUT4 mRNA expression in the LV of male mice at 5 and 12 weeks. Statistical analyses were performed using GraphPad Prism (version 11.0.1). Comparisons between two groups were performed using an unpaired, two-tailed Student’s *t*-test. Statistical significance is indicated as *P* < 0.05, P < 0.01, *P* < 0.001, and P < 0.0001. Abbreviations: GLUT4, glucose transporter 4; RV, right ventricle; LV, left ventricle.

Finally, we assessed PDK4 expression, which is suppressed in the RV of humans with HFpEF[8]. PDK4 inhibits pyruvate dehydrogenase, thus suppressing conversion of the glucose-derived pyruvate to acetyl-CoA. In contrast to observations in humans, our analyses in mice consistently showed a profound upregulation of PDK4 protein and mRNA levels in both sexes, in the RV and LV ventricles, and across disease stages (5 and 12 weeks) (Figure 4A-P).

**Figure 4.**
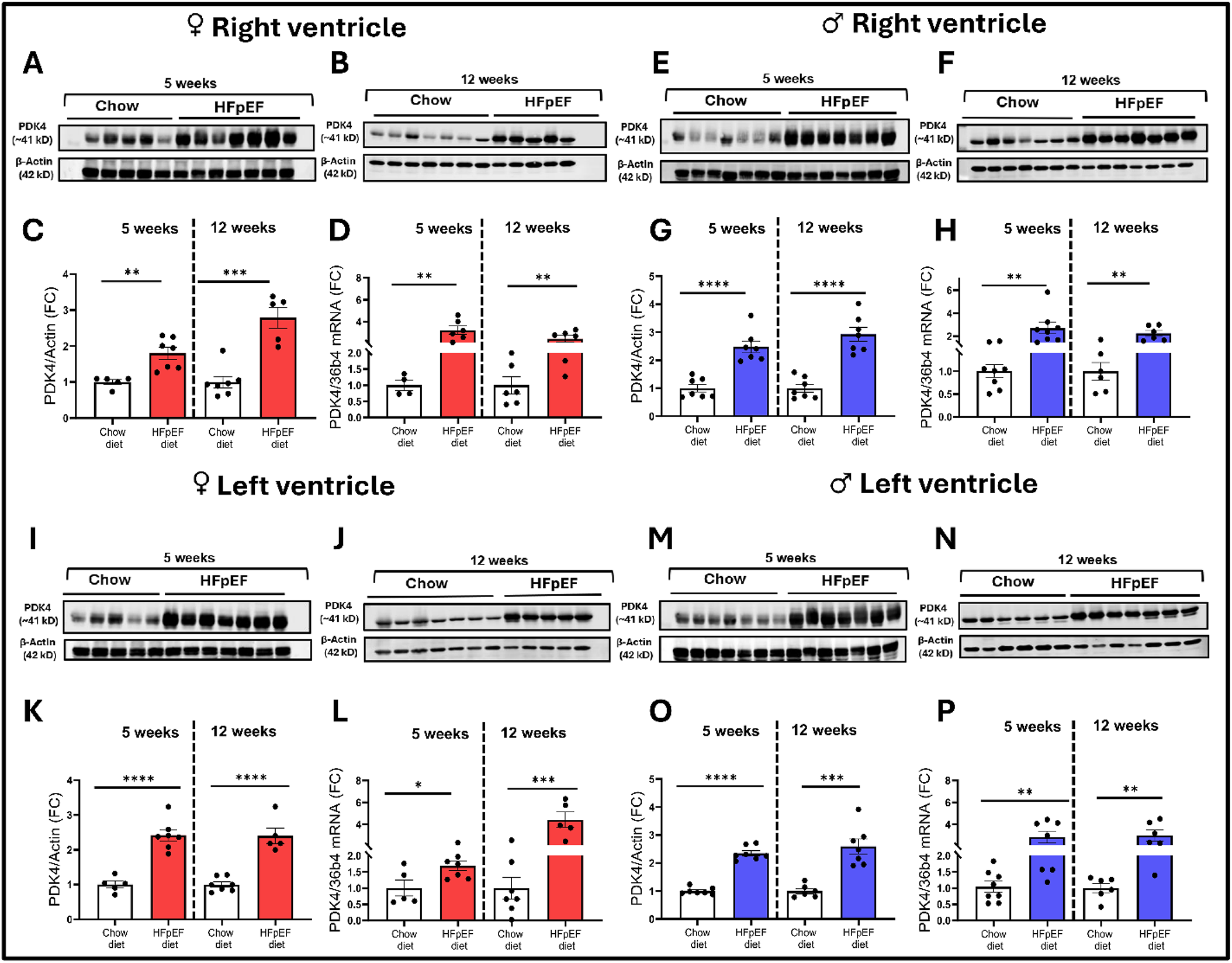
Sex- and chamber-specific regulation of PDK4 in HFpEF. Mice were assigned to either an HFpEF group (high-fat diet plus L-NAME in drinking water; *n* = 5–7 per sex) or a chow-fed control group (*n* = 5–7 per sex). Right ventricular (RV) and left ventricular (LV) tissues were collected after 5 weeks (early phase) and 12 weeks (late phase). PDK4 protein abundance was assessed by immunoblotting with densitometric quantification normalized to β-actin, and PDK4 mRNA expression was normalized to 36B4. (A) Immunoblots of PDK4 protein expression in the RV of female mice at 5 weeks. (B) Immunoblots of PDK4 protein expression in the RV of female mice at 12 weeks. (C) Quantification of PDK4 protein abundance in the RV of female mice at 5 and 12 weeks. (D) PDK4 mRNA expression in the RV of female mice at 5 and 12 weeks. (E) Immunoblots of PDK4 protein expression in the RV of male mice at 5 weeks. (F) Immunoblots of PDK4 protein expression in the RV of male mice at 12 weeks. (G) Quantification of PDK4 protein abundance in the RV of male mice at 5 and 12 weeks. (H) PDK4 mRNA expression in the RV of male mice at 5 and 12 weeks. (I) Immunoblots of PDK4 protein expression in the LV of female mice at 5 weeks. (J) Immunoblots of PDK4 protein expression in the LV of female mice at 12 weeks. (K) Quantification of PDK4 protein abundance in the LV of female mice at 5 and 12 weeks. (L) PDK4 mRNA expression in the LV of female mice at 5 and 12 weeks. (M) Immunoblots of PDK4 protein expression in the LV of male mice at 5 weeks. (N) Immunoblots of PDK4 protein expression in the LV of male mice at 12 weeks. (O) Quantification of PDK4 protein abundance in the LV of male mice at 5 and 12 weeks. (P) PDK4 mRNA expression in the LV of male mice at 5 and 12 weeks. Statistical analyses were performed using GraphPad Prism (version 11.0.1). Comparisons between two groups were performed using an unpaired, two-tailed Student’s *t*-test. Statistical significance is indicated as *P* < 0.05, P < 0.01, *P* < 0.001, and P < 0.0001. Abbreviations: PDK4, pyruvate dehydrogenase kinase 4; RV, right ventricle; LV, left ventricle.

### 3.3 Conserved and Divergent Transcriptional Signatures Across Cardiac Chambers and Sexes in Murine HFpEF

To determine whether sex- and ventricle-specific differences observed in cardiac glucose metabolism-related proteins were reflected at the entire cardiac transcriptome, we performed bulk RNA-seq in the RV and LV of male and female HFpEF and chow-fed control mice (n=5/sex). Principal component analysis (PCA) and sample correlation analysis showed robust segregation of samples by sex, with additional clustering driven by HFpEF status. (Figure 5A, Supplementary Figure 1A). In the RV, female mice with HFpEF exhibited greater dispersion along PC1 and PC2, indicating a more heterogeneous transcriptional response. Differential gene expression analysis (BH-adjusted *P* < 0.05) identified 172 upregulated and 149 downregulated transcripts in female mice and 136 upregulated and 158 downregulated transcripts in males with HFpEF (Figure 5B-C). Hierarchical clustering of the top 30 DEGs clearly separated HFpEF from chow samples in both sexes (Figure 5D-E).

**Figure 5.**
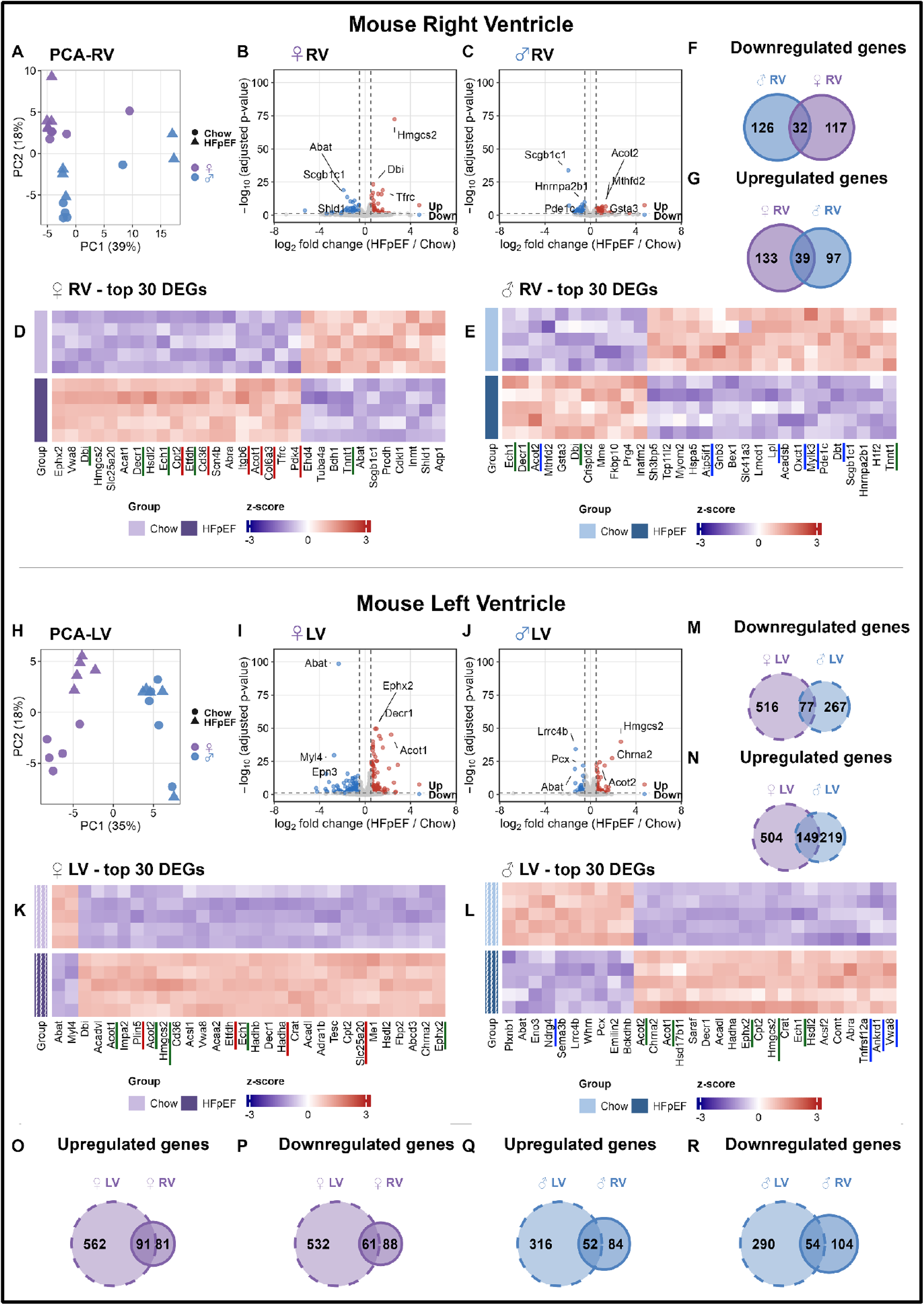
Sex-stratified transcriptomic profiling of right and left ventricles in the mouse HFpEF model. Mice were assigned to either a HFpEF group (high-fat diet plus L-NAME in drinking water; *n* = 5–7 per sex) or a chow-fed control group (*n* = 5–7 per sex). Right ventricular (RV) and left ventricular (LV) tissues were collected after 12 weeks (late phase). Protein-coding genes were retained and filtered for ≥5 counts in ≥25% of samples in either group before differential expression analysis using DESeq2. Genes with a BH-adjusted *P* value < 0.05 were considered differentially expressed. (A) Principal component analysis (PCA) of RV samples. Colors indicate sex (purple, female; blue, male), and shapes indicate treatment (circle, Chow; triangle, HFpEF). (B) Volcano plot of differential gene expression between HFpEF and Chow in the RV of female mice. Upregulated genes are shown in red, downregulated genes in blue, and non-significant genes in gray. The six most significant genes are labeled. (C) Volcano plot of differential gene expression between HFpEF and Chow in the RV of male mice. Upregulated genes are shown in red, downregulated genes in blue, and non-significant genes in gray. The six most significant genes are labeled. (D) Heatmap of the top 30 differentially expressed genes (DEGs), ranked by adjusted *P* value, in the RV of female mice. Values are z-scored by gene, Genes underlined in red are among the top unique DEGs in females, and genes underlined in green are shared between females and males. (E) Heatmap of the top 30 DEGs in the RV of male mice. Values are z-scored by gene. Genes underlined in blue are among the top unique DEGs in males, and genes underlined in green are shared between females and males (F) Venn diagram showing the overlap of downregulated RV DEGs between female and male mice. (G) Venn diagram showing the overlap of upregulated RV DEGs between female and male mice. (H) Principal component analysis (PCA) of LV samples. (I) Volcano plot of differential gene expression between HFpEF and Chow in the LV of female mice. (J) Volcano plot of differential gene expression between HFpEF and Chow in the LV of male mice. (K) Heatmap of the top 30 DEGs in the LV of female mice. Values are z-scored by gene. Genes underlined in red are among the top unique DEGs in females, and genes underlined in green are shared between females and males (L) Heatmap of the top 30 DEGs in the LV of male mice. Values are z-scored by gene. Genes underlined in blue are among the top unique DEGs in males, and genes underlined in green are shared between females and males (M) Venn diagram showing the overlap of downregulated LV DEGs between female and male mice. (N) Venn diagram showing the overlap of upregulated LV DEGs between female and male mice. (O) Cross-chamber Venn diagram comparing upregulated DEGs between the RV and LV of female mice. (P) Cross-chamber Venn diagram comparing downregulated DEGs between the RV and LV of female mice. (Q) Cross-chamber Venn diagram comparing upregulated DEGs between the RV and LV of male mice. (R) Cross-chamber Venn diagram comparing downregulated DEGs between the RV and LV of male mice. Venn diagram areas are proportional to gene-set size. Solid circles indicate RV, and striped circles indicate LV. Abbreviations: HFpEF, heart failure with preserved ejection fraction; PCA, principal component analysis; DEG, differentially expressed gene; RV, right ventricle; LV, left ventricle.

RV transcriptomic changes were highly sex dependent. Both sexes showed increased expression of fatty acid oxidation-related genes, including Ech1 and Decr1. However, RV of female mice exhibited enhanced extracellular matrix remodeling together with increased expression of genes involved in fatty acid uptake and metabolic adaptation, including Col6a3, Itgb6, Hmgcs2, Cd36, and Pdk4, whereas RV of male mice showed reduced expression of genes involved in lipid handling, ketone metabolism, mitochondrial function, and contractile regulation, including *Lpl, Oxct1*, *Dbt*, *Atp5if1*, and *Mylk3*. Consistent with these differences, overlap between sexes was limited, with only 32 commonly downregulated and 39 commonly upregulated shared between sexes (Figure 5F–G).These findings indicate marked sexual dimorphism in RV during HFpEF.

Gene Ontology (GO) and Kyoto Encyclopedia of Genes and Genomes (KEGG) gene sets enrichment analyses further demonstrated marked sex-dependent differences in RV (Supplementary Figure 2). Similar to limited overlap in DEGs, enriched pathways showed minimal concordance between sexes. RV of female mice exhibited activation of inflammatory, extracellular matrix, calcium-handling, and stress-response pathways, together with suppression of contractile and metabolic processes. In contrast, the RV of male mice displayed a more restricted metabolic signature characterized by lipid oxidation, fatty acid metabolism, and adaptive thermogenesis, with limited enrichment in inflammatory pathways.

In the LV, PCA showed a clear separation between chow and HFpEF female mice along PC1 (Figure 5H, Supplementary Figure 3). Compared with the RV, the LV exhibited substantially greater transcriptional alterations. Female mice showed 593 downregulated genes and 653 upregulated, whereas males exhibited 344 downregulated genes and 368 upregulated (Figure 5I-J). Several genes involved in lipid metabolism and mitochondrial function, including *Acot1, Hmgcs2, Acsl1, Ech1, Dbi,* and *Ephx2*, were altered in both sexes (Figure 5K–L). Female LV was further characterized by increased expression of genes involved in fatty acid uptake and oxidation, including *Cd36, Plin5,* and *Slc25a20*, whereas male LV additionally showed differential expression of genes associated with cardiac stress and remodeling, including *Tnfrsf12a, Ankrd1,* and *Ndrg4*.(Figure 5M–N). Pathway analysis of the LV also revealed marked sexual dimorphism (Supplementary Figure 4). Females exhibited strong enrichment of fatty acid oxidation and oxidative phosphorylation-related pathways, together with suppression of multiple immune/cytotoxic pathways. Male mice showed a more limited response, with calcium-ion-regulated exocytosis as the most prominent enriched process.

To identify conserved ventricular responses, DEGs of LV and RV were compared. In females, 91 upregulated and 61 downregulated genes were shared between chambers, indicating that much of the RV transcriptional program mirrors the LV response (Figure 5O-P). In males, 52 upregulated and 54 downregulated transcripts were shared, accounting for of RV DEGs, respectively (Figure 5Q-R). Shared ventricular signatures in both sexes were enriched for fatty acid metabolism, mitochondrial function, and cellular stress responses, including *Hmgcs2, Acadl, Acaa2, Hadha, Hadh, Decr1, Ech1, Cd36, Fabp3, Fabp4, Plin2, Atf5, Nfe2l1, Cat*, and *Ephx2* (Supplementary Table 2). Thus, despite marked sex- and chamber-specific differences, HFpEF is characterized by a conserved core transcriptional program across both ventricles and sexes.

### 3.4. Cross-species cardiac transcriptomics reveals greater conservation of HFpEF-associated gene expression changes in the RV than in the LV

**Reanalysis of** previously published transcriptomic datasets from the RV[14] and LV [13] of HFpEF patients identified robust transcriptional alterations in the RV with relatively few changes in the LV (Figure 6A). To minimize potential confounding cardiovascular HFpEF comorbidities, the datasets were reanalyzed after excluding either patients with AF (Figure 6B), or those with both AF and diabetes (Figure 6C and Supplementary Figure 3). Since these exclusions markedly reduced the sample size, particularly after sex stratification, the male and female datasets were combined for downstream human analyses. The exclusion of patients with AF and diabetes modestly improved cross-species concordance, increasing the number of shared upregulated genes from six to ten and revealing 3 shared downregulated genes, including *CKM, RASD1,* and *CER5* (Figure 6A-C). PCA demonstrated a clear separation between control and HFpEF RV samples (Figure 7A). The human RV exhibited extensive transcriptional reprogramming, with 3,424 upregulated and 3,482 downregulated genes (Figure 7B) with distinct disease-associated clusters (Figure 7C). Cross-species comparison identified 26 commonly upregulated and 43 commonly downregulated genes between human and mouse RV (Figure 7D-E; Supplementary Table 3), enriched in pathways related to mitochondrial function, fatty acid utilization, and cardiac energy metabolism. Key contractile and substrate transport genes (*MYH6, SLC2A1, LPL*) were consistently downregulated, whereas genes involved in lipid and ketone metabolism (*ACAT1, DECR1, ECI2, ACOT2, CIDEA, EPHX2*) were upregulated. These findings suggest that alterations in myocardial energy substrate utilization and mitochondrial metabolism are conserved features of RV HFpEF across species.

**Figure 6.**
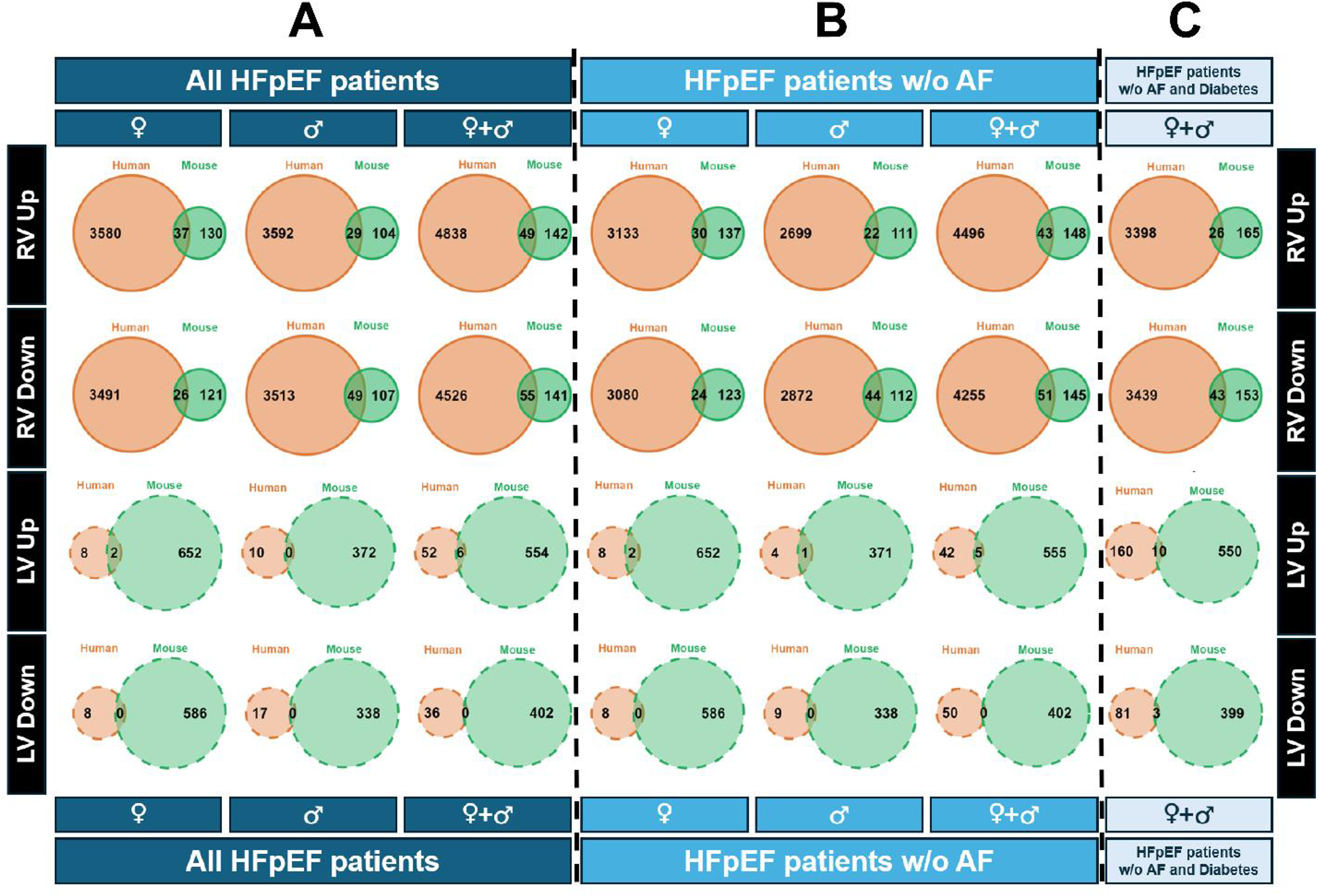
Impact of atrial fibrillation and diabetes exclusion on cross-species transcriptomic concordance in HFpEF. Human HFpEF transcriptomic datasets were analyzed using three filtering strategies: all HFpEF patients, HFpEF patients without atrial fibrillation (AF), and HFpEF patients without AF and diabetes. Protein-coding genes were retained and filtered for ≥5 counts in ≥25% of samples in either group before differential expression analysis using DESeq2. Genes with a BH-adjusted *P* value < 0.05 were considered differentially expressed. Differentially expressed genes (DEGs) identified in human myocardium were compared with DEGs from the HFD/L-NAME murine HFpEF model. Comparisons were performed separately for the right ventricle (RV) and left ventricle (LV) and stratified by sex (female, male, and combined female + male datasets). (A) Venn diagrams showing the overlap of significantly upregulated (Up) and downregulated (Down) DEGs between human and murine HFpEF in the RV and LV using all HFpEF patients. (B) Venn diagrams showing the overlap of significantly upregulated and downregulated DEGs between human and murine HFpEF in the RV and LV after exclusion of patients with AF. (C) Venn diagrams showing the overlap of significantly upregulated and downregulated DEGs between human and murine HFpEF in the RV and LV after exclusion of patients with both AF and diabetes. Numbers indicate the total number of DEGs, and overlapping regions indicate shared genes. Human datasets are shown in orange and murine datasets in green. Solid circles indicate RV, and striped circles indicate LV. Abbreviations: HFpEF, heart failure with preserved ejection fraction; DEG, differentially expressed gene; RV, right ventricle; LV, left ventricle; AF, atrial fibrillation.

**Figure 7.**
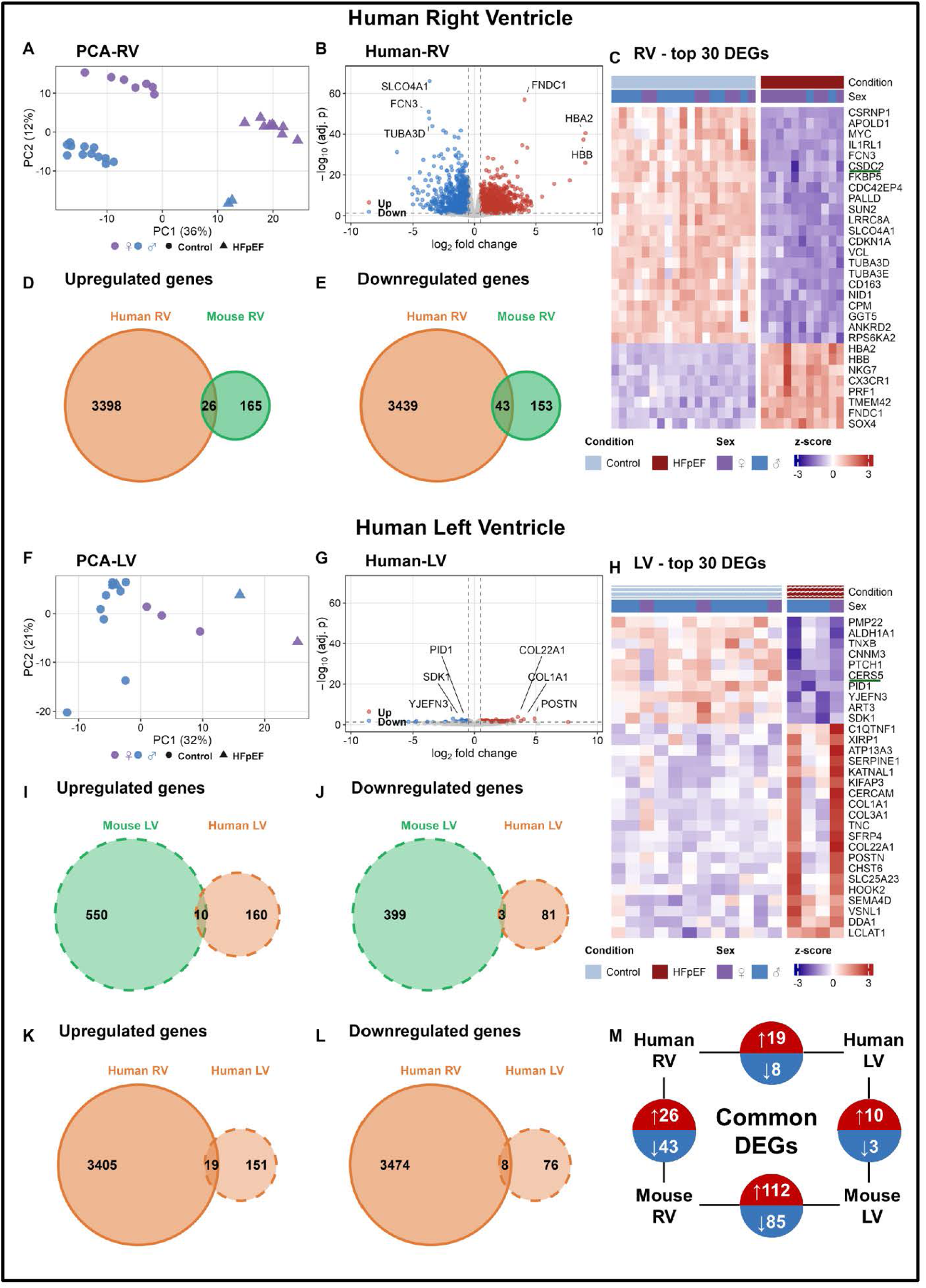
Human HFpEF transcriptomic changes and conservation with the mouse model. Human RV samples were obtained from the RVAD biobank (HFpEF, *n* = 11; Control, *n* = 19), and human LV samples were obtained from the PRJNA879763 dataset (HFpEF, *n* = 4; Control, *n* = 12). Patients with atrial fibrillation or diabetes were excluded from all analyses. Protein-coding genes were retained and filtered for ≥5 counts in ≥25% of samples in either group before differential expression analysis using DESeq2. Genes with a BH-adjusted *P* value < 0.05 were considered differentially expressed. (A) Principal component analysis (PCA) of human RV samples. Colors indicate sex (purple, female; blue, male), and shapes indicate condition (circle, Control; triangle, HFpEF). (B) Volcano plot of differential gene expression between HFpEF and Control in the human RV. Upregulated genes are shown in red, downregulated genes in blue, and non-significant genes in gray. The six most significant genes are labeled. (C) Heatmap of the top 30 differentially expressed genes (DEGs), ranked by adjusted *P* value, in the human RV. Values are z-scored by gene, green underline represents the common gene between human and mouse. (D) Venn diagram showing the overlap of upregulated DEGs between the human and mouse RV. (E) Venn diagram showing the overlap of downregulated DEGs between the human and mouse RV. (F) Principal component analysis (PCA) of human LV samples. (G) Volcano plot of differential gene expression between HFpEF and Control in the human LV. (H) Heatmap of the top 30 DEGs, ranked by adjusted *P* value, in the human LV, genes underlined in green are shared between the human and mouse datasets. Values are z-scored by gene. (I) Venn diagram showing the overlap of downregulated DEGs between the human and mouse LV. (J) Venn diagram showing the overlap of upregulated DEGs between the human and mouse LV. (K) Venn diagram showing the overlap of upregulated DEGs between the human RV and LV. (L) Venn diagram showing the overlap of downregulated DEGs between the human RV and LV. (M) Summary of common genes identified across human and mouse ventricles. Venn diagram areas are proportional to gene-set size. Human gene symbols were matched to mouse orthologs for all cross-species comparisons. Abbreviations: HFpEF, heart failure with preserved ejection fraction; PCA, principal component analysis; DEG, differentially expressed gene; RV, right ventricle; LV, left ventricle.

In comparison, PCA of the human LV dataset showed substantial overlap between control and HFpEF samples, indicating less distinct transcriptional patterns and greater inter-sample heterogeneity (Figure 7F). The LV also exhibited limited transcriptional changes, with 84 downregulated and 170 upregulated genes (Figure 7G). Unlike the RV, the LV did not display a strongly segregated disease signature. However, heatmap analysis still demonstrates robust transcriptomics changes in HFpEF patients compared to healthy groups (Figure 7H). Similarly, comparison of the LV datasets between humans and mice with HFpEF identified ten common upregulated (Figure 7I) and only three common downregulated genes (Figure 7J) between human and mouse samples. Shared downregulated genes, including *CKM, RSAD1*, and *CERS5*, suggest impaired energy buffering and altered lipid signaling. Conversely, conserved upregulated genes such as *HSPA1A, ITGB1, ADAMTS9,* and *FKBP10* point to enhanced protein quality control and extracellular matrix reorganization (Supplementary Table 4). As with the murine model, comparison of the human RV and LV revealed only 19 common upregulated transcripts (Figure 7K), and eight common downregulated transcripts (Figure 7L), highlighting even less -compared to mice-overlapping transcriptional signatures among the two ventricles differences. Similarly, Comparison of GO and KEGG enrichment analyses between mouse and human HFpEF datasets revealed predominantly immune, inflammatory, and cell cycle-related signatures. Notably, shared enrichment patterns were more extensive in the RV than in the LV, indicating greater cross-species similarity in RV (Supplementary Figure 5,6).

Collectively, these findings identify conserved biological pathways despite limited transcript overlaps, with greater cross-species concordance in the RV than the LV (Figure 7M).

## 4. Discussion

HFpEF accounts for more than 50% of all HF cases, yet effective therapies remain lacking[16, 17]. Women are disproportionately affected[18, 19], underscoring a critical gap in our understanding of the molecular mechanisms driving sex-specific vulnerability in HFpEF. Our study aims to identify patterns of either universal or chamber-specific transcription reprograming among mice and humans, as well common and divergent gene expression patterns in males and females. Our findings reveal a striking misalignment in GLUT1 mRNA and protein levels in the RV of male mice, which mirrors observations that have been reported in human HFpEF RV tissue[8]. Furthermore, we identified conserved transcriptomic signatures involved in cardiac bioenergetics, as well as chamber- and sex-specific molecular targets shared between human HFpEF and the “two-hit” mouse model, highlighting biologically relevant target with strong translational potential across species.

Glucose uptake in the heart is primarily mediated by GLUT1 and GLUT4 [4]. The physiological preference in fatty acid oxidation for myocardial ATP production [20] deviates in the stressed heart, when glucose utilization typically increases[21]. However, in HFpEF, this reprogramming does not restore energetic homeostasis[22] as HFpEF hearts exhibit a more uncoupled metabolic phenotype[8, 9, 22]. In the RV of humans with HFpEF, GLUT1 protein levels are increased while dissociation between glucose uptake and downstream oxidative metabolism occurs[8, 22]. Our observations in the RV of male mice with HFpEF are consistent with these findings. Thus, imbalanced glucose handling in the RV of male mice may be an important feature of HFpEF. Notably, clinical trials including EMPEROR-Preserved[6], DELIVER, STEP-HFpEF DM [7], and SUMMIT[23] have shown that glucose lowering treatments, such as SGLT2 inhibitors and GLP-1 receptor agonists, improve cardiovascular outcomes in HFpEF [6, 7]. In contrast, GLUT1 in the LV was reduced at both the mRNA and protein levels in male and female mice, while no change in transcript levels was observed in human LV. Recent human LV proteomic and metabolomic studies[22] have, likewise, reported reduced glycolytic and TCA intermediates, supporting the concept that altered myocardial glucose utilization is a key feature of HFpEF and warranting further mechanistic study of the LV.

Finally, a similar divergence was observed for PDK4, increased in mice and reduced in previously published human HFpEF. Given the frequent coexistence of HFpEF with comorbidities in humans, it remains unclear whether these differences reflect disease stage-dependent effects or the influence of patient medications.

The RV and LV differ markedly in structure and function. The LV has a thicker myocardium adapted for high-pressure systemic circulation, whereas the RV is specialized for low-pressure pulmonary circulation[24, 25]. Although LV dysfunction is often emphasized as a hallmark of HFpEF, increasing evidence indicates that RV dysfunction also contributes substantially to morbidity and mortality in HF [26]. Given these intrinsic differences, molecular changes observed in the LV cannot be assumed to occur similarly in the RV[22, 25]. Indeed, our ventricular-specific transcriptomic profiling in murine revealed a striking chamber-dependent pattern of transcriptome changes, with only modest overlap between LV and RV gene sets that does not exceed 20% of DEGs in both females and males. Strikingly similar observation was observed when comparing the human RV and LV. These findings suggest that LV and RV undergo largely distinct transcriptional alteration programs, with only a limited common gene set likely reflecting shared cardiac stress responses. This finding highlights the importance of considering RV as a biologically distinct compartment rather than an extension of LV pathology. This discrepancy needs to be taken into account when systemic pharmacologic treatments are designed[27]. On the other hand, the conserved genes shared between chambers, many of which are involved in metabolic adaptation, mitochondrial function, and cellular stress responses, may represent attractive therapeutic targets that modulate pathological alteration in both ventricles.

Although HFpEF affects both sexes, females show higher risk of morbidity and mortality[28]. Postmenopausal women, in particular, exhibit higher incidences of ventricular stiffness, fibrosis, and diastolic dysfunction, leading to poorer clinical outcomes[29, 30]. Our findings highlight a distinct sexual dimorphism in HFpEF. While male mice exhibited relatively limited transcriptional changes in both ventricles, female mice demonstrated extensive chamber-specific alteration. The RV of female mice was characterized by enrichment of immune and inflammatory pathways, including leukocyte activation, T-cell signaling, and cytokine production. In contrast, the LV of female mice exhibited enrichment of fatty acid metabolism, mitochondrial respiration, and electron transport chain pathways, together with reduced immune signaling. These findings suggest that inflammation predominates in the RV, whereas metabolic alteration predominates in the LV of female HFpEF mice. This is consistent with prior work showing that sex differences in cardiac mitochondrial function and fatty acid metabolism contribute to diastolic dysfunction and HFpEF susceptibility in female mice compared to male[31].

Lastly, the use of high-fidelity animal models that accurately replicate HFpEF metabolic phenotype and can be readily subjected to pharmacologic or genetic interventions is essential for deciphering HFpEF pathophysiology and identifying novel therapeutic targets. Several mouse models recapitulate features of HFpEF, including aging C57BL/6 mice[32, 33], senescence-accelerated mice[34], hyperphagia models such as ob/ob[35] and db/db mice[36], the SAUNA model[37], and models driven by angiotensin II infusion[38] or mild transverse aortic constriction combined with deoxycorticosterone treatment[39]. The “two-hit” mouse model, which was employed in the present study, constitutes the most widely used pre-clinical model for HFpEF[12]. This model closely recapitulates key features of human HFpEF and offers several practical advantages, including non-surgical induction through dietary and drinking water modifications, a relatively short treatment duration (5–15 weeks), and reproducible diastolic dysfunction with preserved systolic function across various genetic backgrounds, such as C57BL/6N, C57BL/6J, and FVB/N[12, 40, 41]. The discrepancies between human HFpEF and mouse models may reflect species-specific disease mechanisms or the influence of comorbidities, including diabetes, metabolic syndrome, atrial fibrillation, and hypertension. Better clinical characterization of HFpEF populations will improve the translatability of the “two-hit” and other preclinical models.

Despite the limited overlap between murine and human HFpEF transcriptomes, the RV exhibited a conserved molecular signature centered on metabolic alteration and mitochondrial dysfunction in both species. Shared transcriptome changes included upregulation of genes involved in mitochondrial fatty acid metabolism and energy substrate utilization, alongside lower expression of genes associated with glucose transport and cardiac energy metabolism. Given that RV dysfunction is increasingly recognized as an independent predictor of mortality and HF hospitalization in HFpEF[42], the identification of conserved metabolic alterations in the RV have valuable translational implications for understanding RV pathophysiology and developing targeted therapeutic strategies. In contrast, only limited overlap was observed in the LV. Nevertheless, the common transcript changes identified in the LV of humans and mice were associated with pathways involved in cellular stress responses, protein homeostasis, and structural changes, which are important for HFpEF [43, 44]. For example, Upregulation of *HSPA4* and *CPEB4* suggests adaptive stress responses and protein quality control, whereas increased *ITGB1, ADAMTS9, and DYSF* is consistent with mechanotransduction, extracellular matrix remodeling, and membrane repair. Downregulation of *CKM* and *CERS5* suggests impaired energy metabolism and altered lipid signaling.

## Conclusion

In summary, our comparative cardiac transcriptomic and targeted protein analyses demonstrate that HFpEF is characterized by chamber-specific, sex-dependent, and species-dependent metabolic alteration. The RV exhibited the most robust and conserved disease-associated signature compared to LV. Notably, female mice exhibited distinct chamber-specific responses, with inflammatory changes predominating in the RV and metabolic alteration in the LV. Although key regulators such as *GLUT1, PDK4*, and *GLUT4* were only partially conserved between human HFpEF and the HFD/L-NAME mouse model, the shared alterations converge on a focused set of metabolic pathways. These conserved molecular changes provide strong candidates for future mechanistic studies aiming to translate across species.

## 5. Limitation and future perspective

Several limitations should be considered when interpreting these findings. First, the relatively small sample size and unequal sex distribution in the human LV cohort constitute important limitations, potentially reducing statistical power. In addition, male and female mice were studied on different C57BL/6 strains, as male mice were C57BL/6N and female mice were C57BL/6J because it was previously suggested that female C57BL/6N mice are resistant in HFpEF development [45]. Although we excluded patients with atrial fibrillation and diabetes, the human datasets likely still reflect residual clinical heterogeneity due to additional cardiovascular comorbidities, disease stage and medication effects.

Our analysis was restricted to protein-coding genes and bulk tissue, which limits the detection of non-coding regulatory RNAs and cell type-specific altered gene expression profile. Future single-cell and spatial transcriptomic studies will be important for defining the cellular and regional origins of these changes. The human HFpEF datasets reanalyzed in this study were generated by independent research groups using different patient cohorts and sampling time points, which may partially explain the marked disparity in the number of transcriptomic alterations observed between the RV and LV. Although the number of DEGs in the LV increased following the exclusion of patients with comorbidities such as AF and diabetes, these findings should be interpreted with caution, as residual comorbidities and methodological heterogeneity may still influence the observed transcriptional signatures.

## CREDIT AUTHORSHIP CONTRIBUTION STATEMENT

K. Thapa: Writing – Original draft, Methodology, Formal analysis, Data curation. KM Verrou-Methodology, Formal analysis, Data curation. E. Rapushi: Methodology, Data curation. G. Siokatas – Methodology, Data curation. N. Bharucha – Methodology. K Chella Krishna-Writing review and editing. B. J Keating-review and editing, M Meyer-review and editing, I Karakikes-review and editing, K. Drosatos – Writing, review and editing, Supervision, Resources, Project administration, Investigation, Funding acquisition, Conceptualization.

---

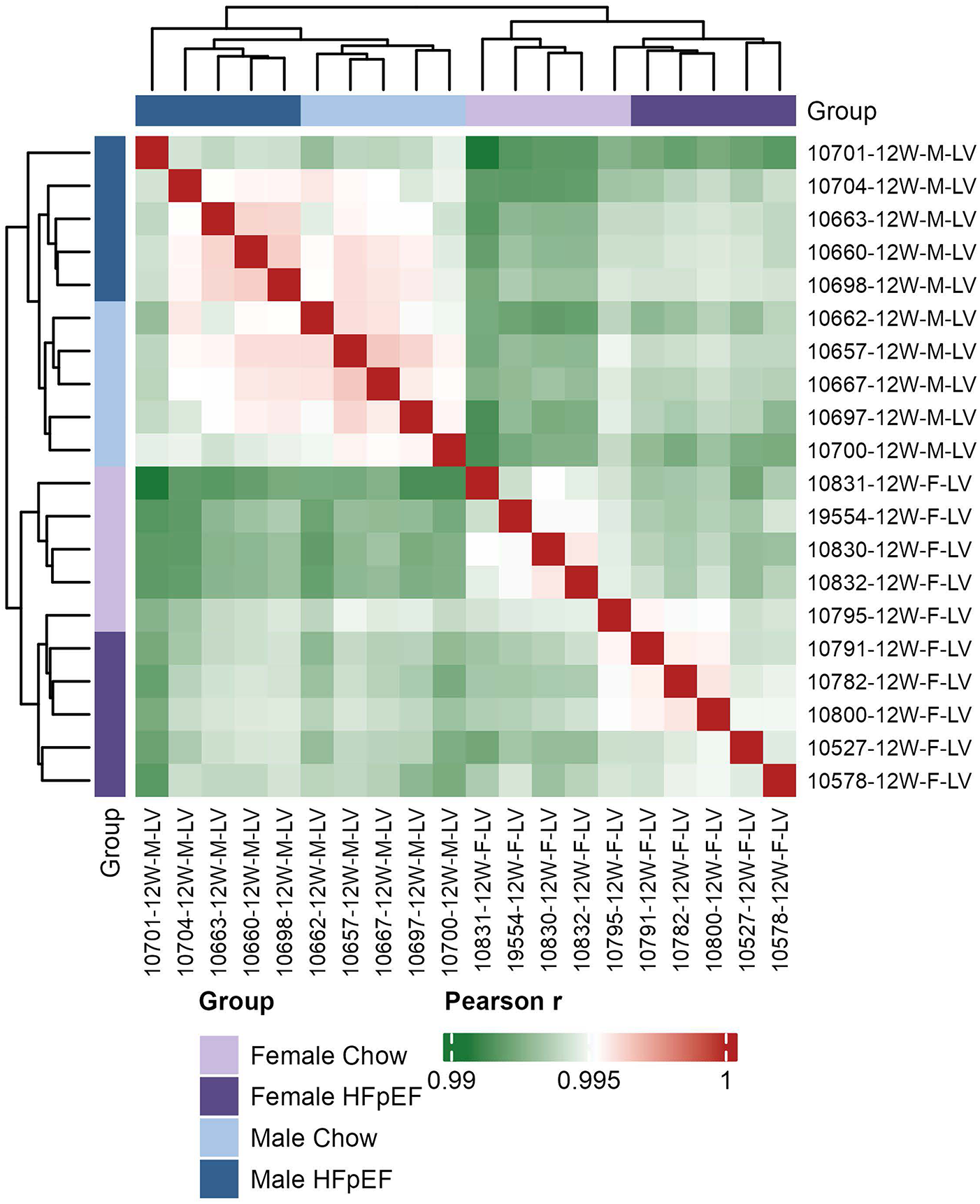

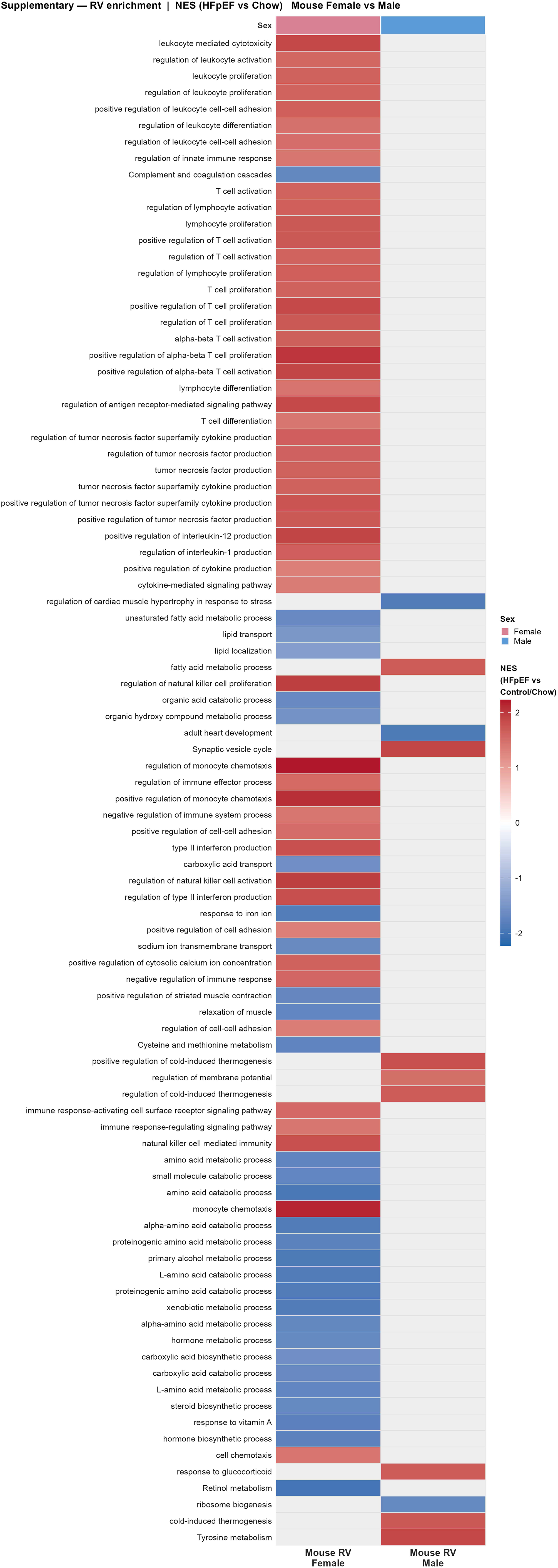

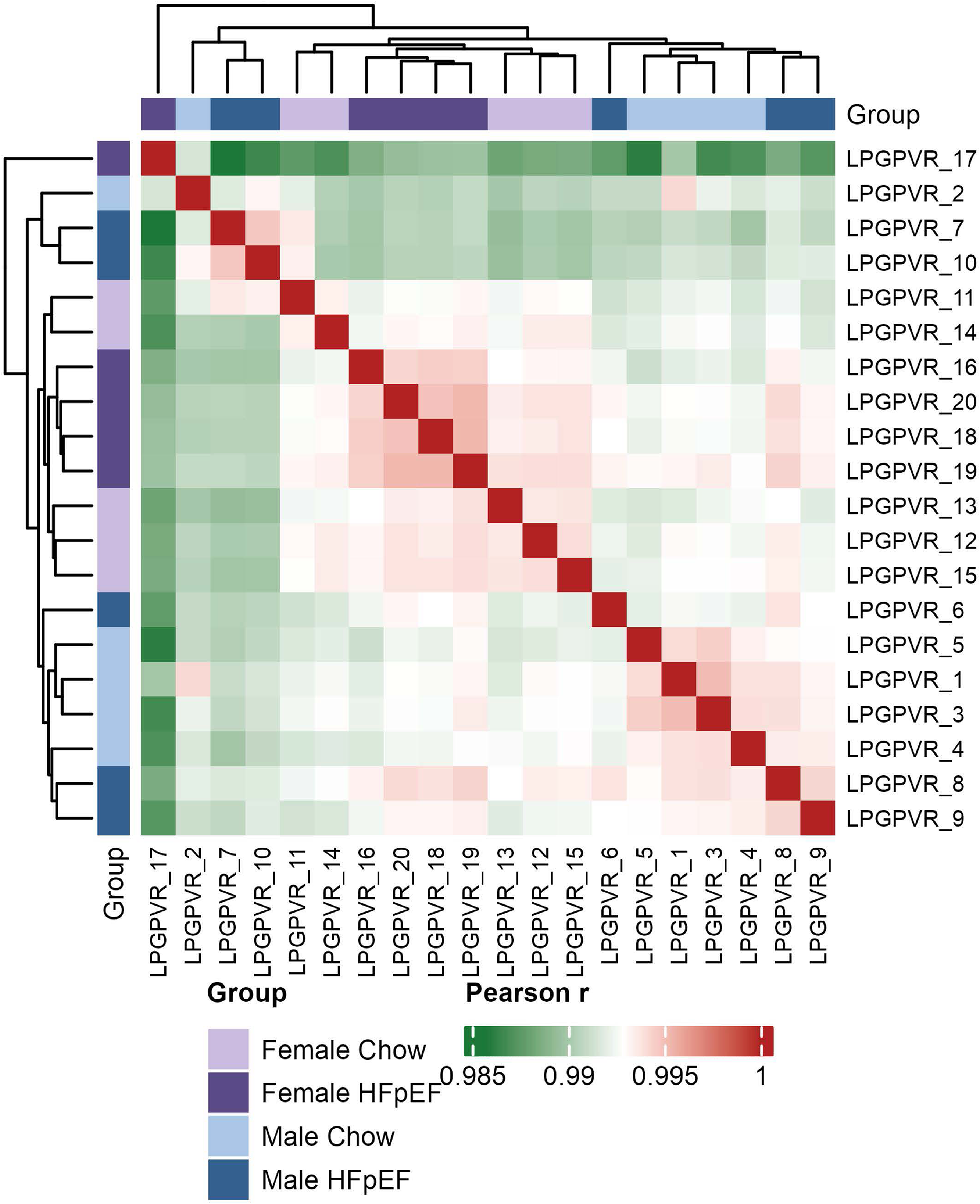

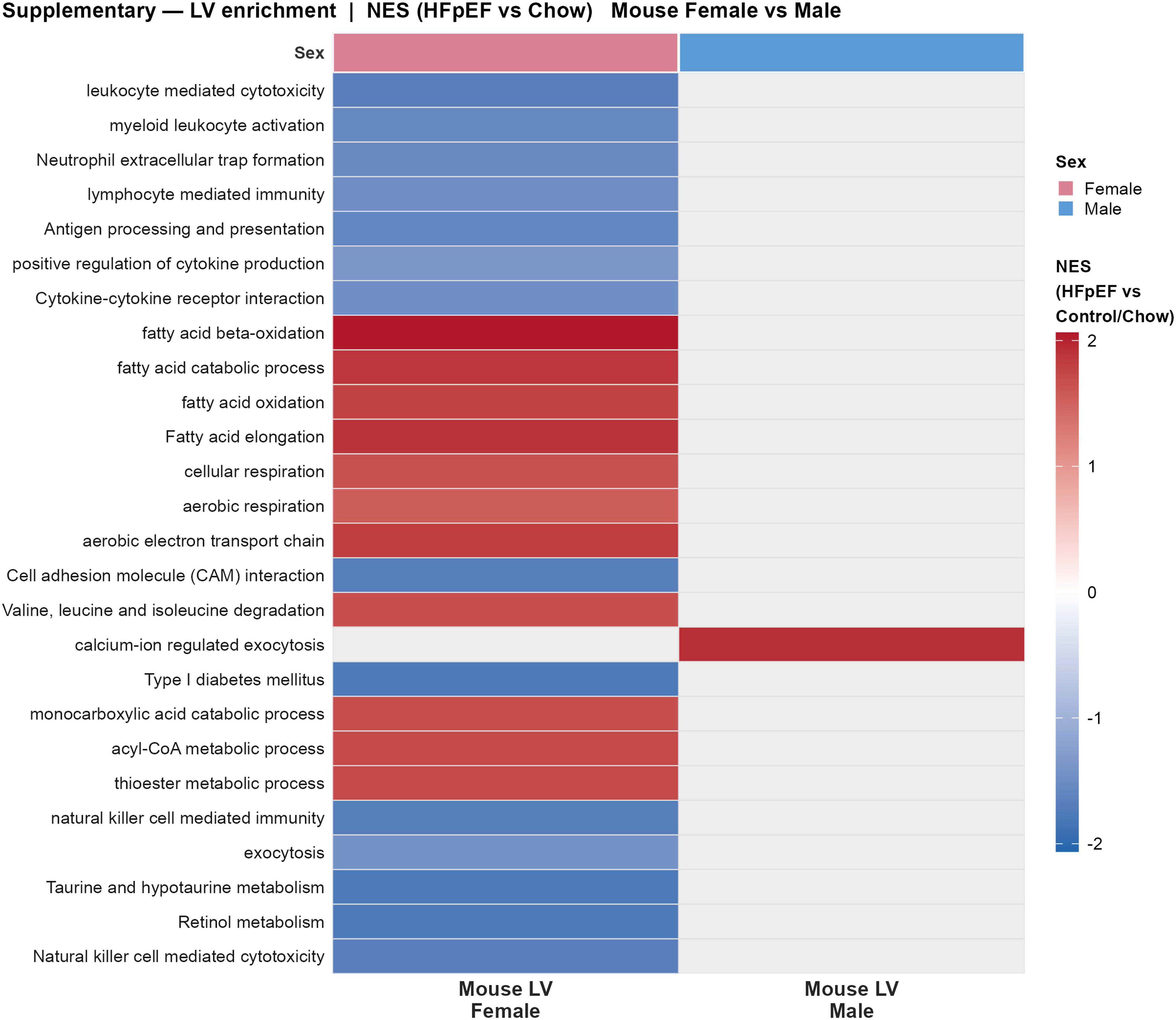

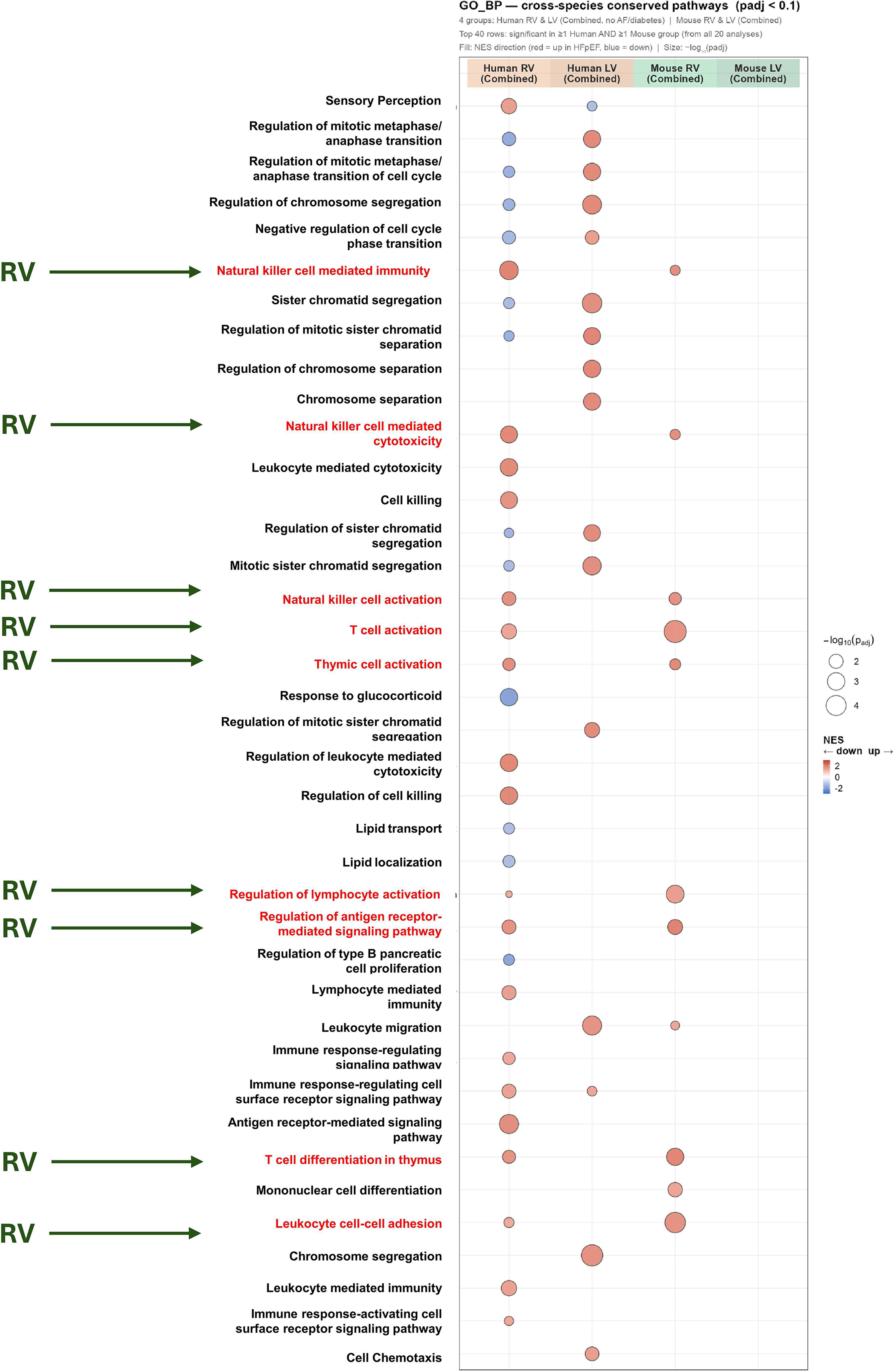

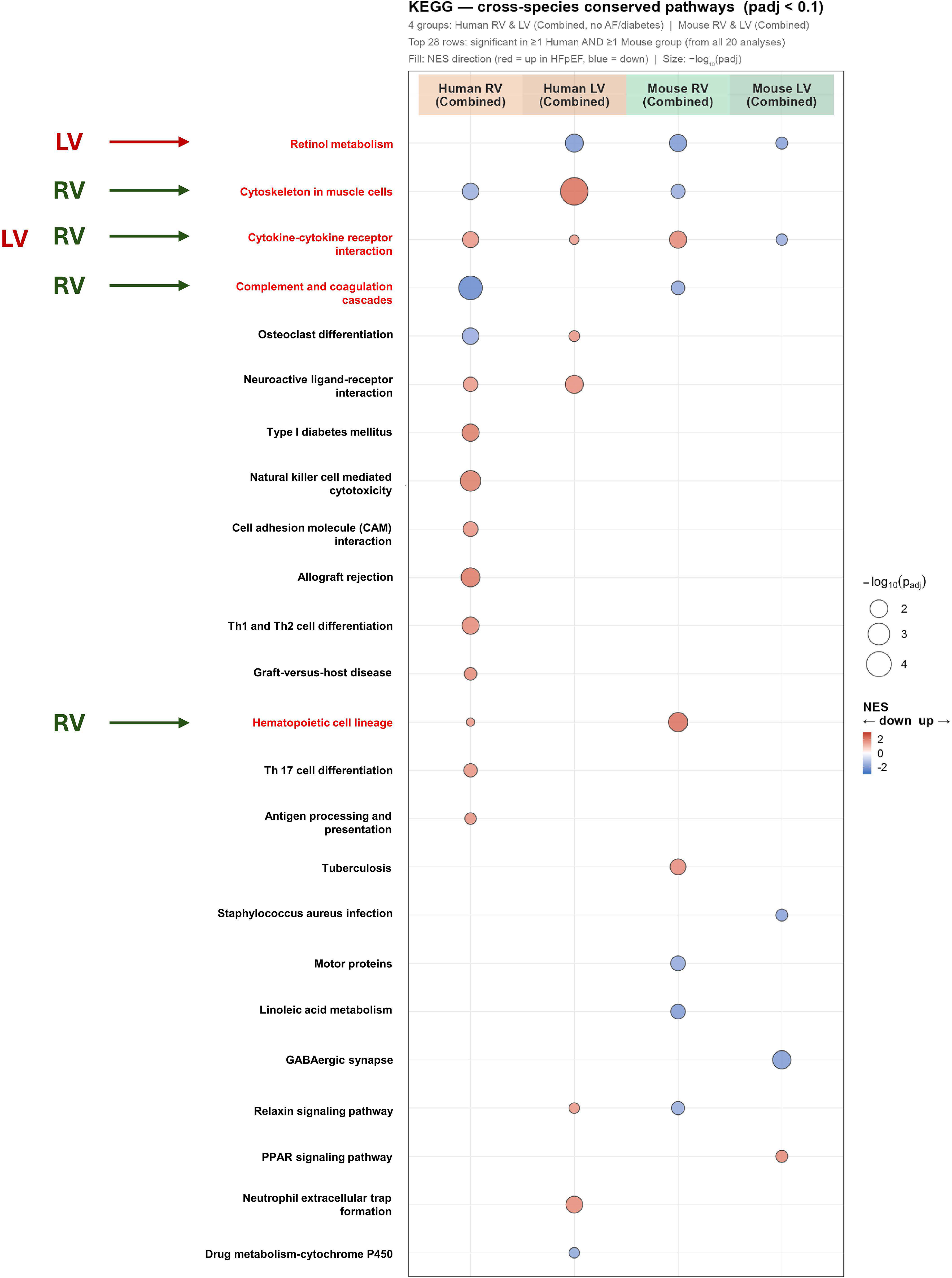

